# Decomposing the admixture statistic, D, suggests a negligible contribution due to archaic introgression into humans

**DOI:** 10.1101/2021.01.21.427635

**Authors:** William Amos

**Author notes:** Author and corresponding author: William Amos.

## Abstract

It is widely accepted that non-African humans carry a few percent of Neanderthal DNA due to historical inter-breeding. However, methods used to infer a legacy all assume that mutation rate is constant and that back-mutations can be ignored. Here I decompose the widely used admixture statistic, D, in a way that allows the overall signal to be apportioned to different classes of contributing site. I explore three main characteristics: whether the putative Neanderthal allele is likely derived or ancestral; whether an allele is fixed in one of the two human populations; and the type of mutation that created the polymorphism, defined by the base that mutated and immediately flanking bases. The entire signal used to infer introgression can be attributed to a subset of sites where the putative Neanderthal base is common in Africans and fixed in non-Africans. Moreover, the four triplets containing highly mutable CpG motifs alone contribute 29%. In contrast, sites expected to dominate the signal if introgression has occurred, where the putative Neanderthal allele is absent from Africa and rare outside Africa, contribute negligibly. Together, these observations show that D does not capture a signal due to introgression but instead they support an alternative model in which a higher mutation rate in Africa drives increased divergence from the ancestral state.

## Introduction

Over the last decade, the revelation that humans inter-bred with Neanderthals and other archaic hominins, leaving appreciable genetic legacies in all non-Africans, has transitioned from surprising inference to textbook fact (1–4). During this time, a remarkable weight of evidence has accumulated, represented by literally hundreds of publications. We now understand that while some introgressed fragments were selected against (5, 6), others appear to have facilitated adaptation (7–9). Moreover, introgression seems to be prevalent among related taxa such as chimpanzees (10). Widespread introgression is somewhat puzzling both behaviourally, because diverse behavioural mechanisms usually discourage inter-specific mating (11), and evolutionarily, because gene flow tends to prevent speciation.

Most inferences concerning introgression rely either directly or indirectly on the twin assumptions that mutation rate is constant and that back-mutations are too rare to have an appreciable impact (1, 12, 13). Under these assumptions, archaics will share the same number of bases with all human populations unless base-sharing is increased by introgression. The tendency of archaics to share more bases with one human population compared with a second is often quantified with the statistic, D (1, 13) and related measures (14). However, a closer look at D reveals patterns that are not expected under the inter-breeding model. Crucially, inter-breeding predicts that most introgressed bases will be rare, and will therefore tend be found in the heterozygous state in non-Africans, yet D is actually dominated by sites where the African individual is heterozygous (15).

The only alternative to the inter-breeding hypothesis is appears to be a model based on variation in mutation rate. Specifically, the observed tendency for Neanderthals to share more bases with non-Africans compared with Africans could be driven by a higher mutation rate in Africa, driving higher divergence from the ancestral state. Mechanistically, evidence is accumulating that mutation rate is higher at and around heterozygous sites (16–18). If so, then the large loss of heterozygosity seen in humans who left Africa to colonise the rest of the world would have caused a parallel reduction in mutation rate. Such a reduction has been observed: genomic regions that lost most heterozygosity out of Africa also show the largest excess mutation rate in Africa (18). Elsewhere, Africans and non-Africans exhibit striking differences in their mutation spectra (19, 20), showing that something about the mutation process is changing on the timescale of global human expansion. Tying these observations together, the relative mutation rate of different base combinations, *sensu* Harris (19), is impacted by flanking sequence heterozygosity (21), with some base combinations increasing in relative probability in more heterozygous regions and others decreasing.

Here I conduct a further test of the idea that mutation rate variation drives D, rather than genuine introgression. I do this by partitioning D according to the mutation spectrum, testing the prediction that large D values will be associated preferentially with sites that have inherently higher mutation rates.

## Results

### a) Decomposing D by type of site

D is typically calculated from four-way alignments comprising two humans, a test group and an outgroup, written as D(*P1, P2, P3, O*) (13, 22, 23). In the now classic comparison, the two humans were drawn from *P1* = Europe and *P2* = Africa, the test group *P3* = Neanderthal and *O* = chimpanzee. Informative bases are usually taken as bi-allelic sites where the chimpanzee base (always labelled ‘A’) and Neanderthal base (always ‘B’) differ. Where humans carry both bases, there are two possible states, ABBA and BABA. D is calculated as the normalised difference in counts of these two states: (BABA – ABBA)/(BABA + ABBA), where BABA and ABBA are the numbers of each type found across the genome, either directly scored in two individual humans, or probabilistically based on allele frequencies in two population samples. However, if the BABA – ABBA difference is not normalised, it can be decomposed simply by counting ABBAs and BABAs attributable to different classes of site.

Following Harris and colleagues (19, 20), I classify substitutions according to the base that mutates and the two immediately flanking bases, giving a total of 64 triplets. However, each triplet on one strand has an equivalent on the other strand, allowing the full 64 triplets to be collapsed to 32 triplets with either A or C as the central base. I focused on three representative population comparisons: non-African – African (*P1* = GBR, *P2* = ESN); one extreme non-African – non-African comparison between populations at opposite ends of Eurasia (*P1* = GBR, *P2* = CHB); and one within Africa comparison (*P1* = LWK, *P2* = ESN). Within each comparison I also classified sites according to whether the Neanderthal B allele was likely ancestral (Neanderthal = Denisovan) or derived (Neanderthal ≠ Denisovan), yielding the six comparison – site type combinations. Summary counts not broken down by triplet are presented in Table 1, while Figure 1 presents these same data broken down by triplet. Comparisons based on other population combinations yield plots that conform closely to type (i.e. all non-African – African comparisons are similar) and are presented as regional averages in ESM2.

**Table 1.** Summary counts across all triplets. BABA – ABBA counts are generated according to the type of site involved: (i) status of the Neanderthal B allele, likely ancestral (ANC) or derived (DVD) and both ANC + DVD together (ALL); (ii) if and which allele is fixed in one of the two population, e.g. A_P1 indicates that the A allele is fixed in population *P1* (and hence segregating in P2), POLY = segregating in both populations; (iii) population comparison, illustrated by Europe-Africa (GBR-ESN), Europe-East Asia (GBR-CHB) and two African populations (LWK-ESN). Counts are presented both as total counts, BABA+ABBA (upper half) and as BABA-ABBA differences (lower half). D can be obtained by diving any given value in the lower half of the table by the equivalent value in the upper half.

|  |  |  | ALL | B_P1 | A_P1 | B_P2 | A_P2 | POLY |
| --- | --- | --- | --- | --- | --- | --- | --- | --- |
| BABA + ABBA | ALL | EUR-AFR | 320089 | 27922 | 11359 | 2850 | 8353 | 269606 |
|  |  | EUR-EAS | 257586 | 3056 | 6190 | 4702 | 5683 | 237949 |
|  |  | AFR-AFR | 318258 | 931 | 176 | 1487 | 553 | 315114 |
|  | ANC | EUR-AFR | 223100 | 23731 | 6106 | 2751 | 2106 | 188406 |
|  |  | EUR-EAS | 173591 | 2696 | 2138 | 4102 | 2152 | 162496 |
|  |  | AFR-AFR | 226357 | 901 | 71 | 1424 | 203 | 223762 |
|  | DVD | EUR-AFR | 96989 | 4192 | 5253 | 99 | 6247 | 81200 |
|  |  | EUR-EAS | 83995 | 360 | 4051 | 600 | 3531 | 75453 |
|  |  | AFR-AFR | 91900 | 30 | 105 | 64 | 350 | 91351 |
| BABA - ABBA | ALL | EUR-AFR | 24807 | 27922 | -11359 | -2850 | 8353 | 2745 |
|  |  | EUR-EAS | -1962 | 3056 | -6190 | -4702 | 5683 | 191 |
|  |  | AFR-AFR | 416 | 931 | -176 | -1487 | 553 | 595 |
|  | ANC | EUR-AFR | 16296 | 23731 | -6106 | -2751 | 2106 | -679 |
|  |  | EUR-EAS | -577 | 2696 | -2138 | -4102 | 2152 | 815 |
|  |  | AFR-AFR | 217 | 901 | -71 | -1424 | 203 | 607 |
|  | DVD | EUR-AFR | 8511 | 4192 | -5253 | -99 | 6247 | 3424 |
|  |  | EUR-EAS | -1385 | 360 | -4051 | -600 | 3531 | -625 |
|  |  | AFR-AFR | 199 | 30 | -105 | -64 | 350 | -12 |

**Figure 1.**
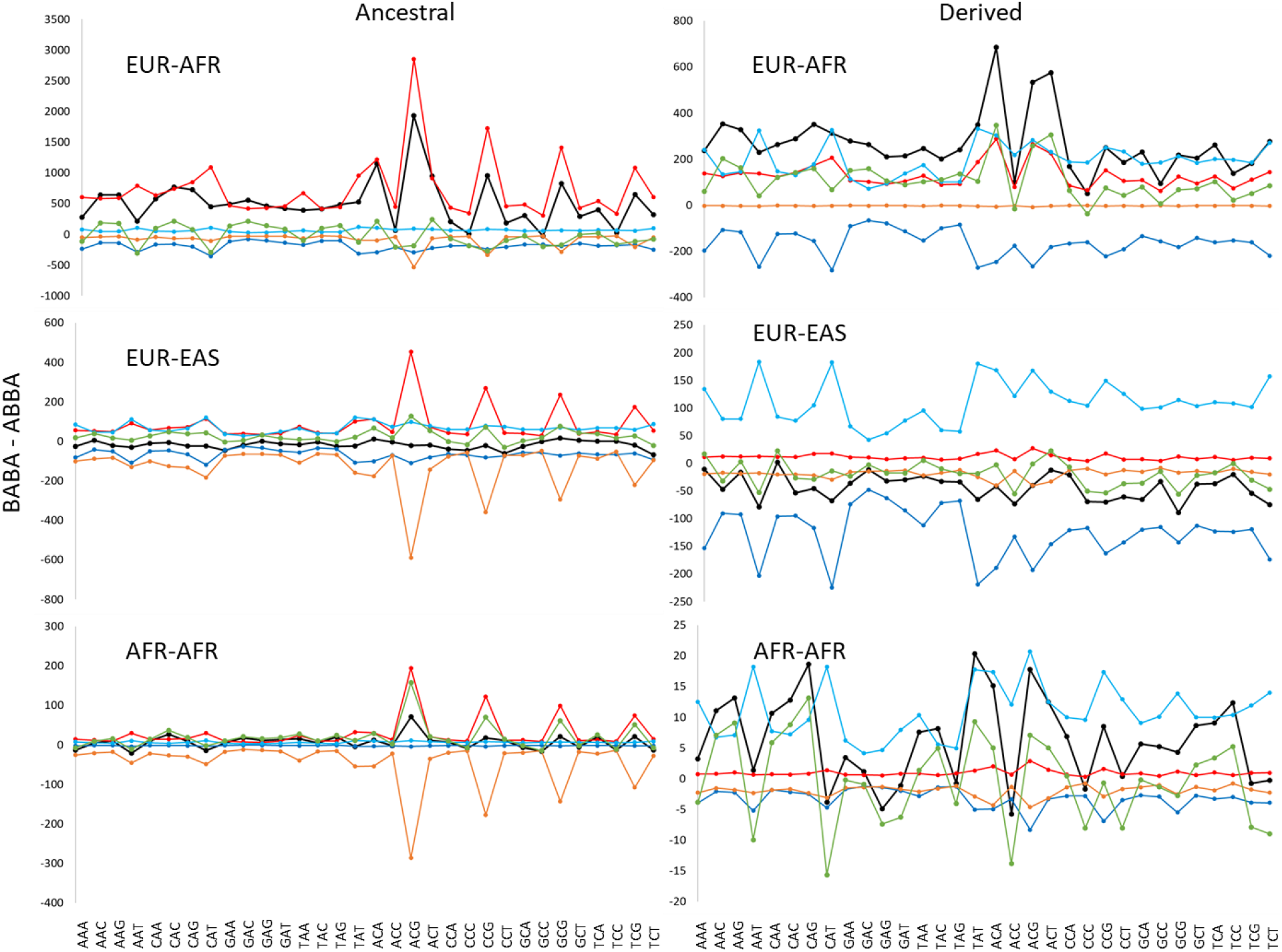
Decomposition of D according to the types of site that contribute. Every site contributing to the classic admixture statistic D(*P1,P2,Necnderthcl,chimpcnzee*) was classified according to (i) the type of base that mutated to create the substitution, sensu Harris (19), defining 32 different mutating triplets (X-axis); (ii) whether the Neanderthal allele was likely ancestral (Neanderthal = Denisovan, left-hand panels) or derived (Neanderthal Denisovan, right-hand panels); and (iii) according to three representative pairwise population comparisons [Europe – Africa (GBR – ESN, top panels), Europe – East Asia (GBR – CHB, middle panels), Africa – Africa (LWK – ESN, bottom panels)]. Within each of the resulting six panels, BABA – ABBA counts are presented either as totals (black) or as sub-counts based on sites where: *P1* is fixed for B (red); *P1* is fixed for A (orange); *P2* is fixed for B (dark blue); *P2* is fixed for A (light blue); A and B present in both human populations (green). Lines between points are added to aid clarity and have no other meaning. In all cases, positive values indicate *P1* shares more bases than *P2* with the Neanderthal.

Figure 1 comprises six panels. In each panel, the X-axis divides the data into 32 triplets while the Y-axis is total BABA-ABBA count differences for all D-informative sites (black) plus five sub-classes of site where: B is fixed in *P1* (red); B is fixed *P2* (orange); A is fixed in *P1* (dark blue); A is fixed in *P2* (light blue); both alleles present in *P1* and *P2* (green). Without any decomposition, i.e. the value used to calculate classical D, BABA – ABBA = 24,807. I refer to this as the total difference and it is similar to but smaller than the total difference contributed by sites where *P1* is fixed for B (27,922) and only marginally larger than the number contributed by sites where the B allele is both fixed in *P1* and likely ancestral (23,731). Considering first the top left panel (Europe – Africa, ancestral B alleles), four triplets dominate the profile: ACG, CCG, GCG and TCG, together accounting for a BABA – ABBA difference of 7,074, 28.5% of the unpartitioned total. These are the four triplets where the central base is the C of a CpG, known to be highly mutable (24–26). Moreover, there is a close match between the overall difference (black) and differences at sites where the B allele is ancestral and fixed in Europe (red), the overall counts tending to be lower, particularly for the four CpG triplets.

Moving to consider ancestral B alleles in the two within-region comparisons (middle left and bottom left panels), related profiles are seen, with CpG triplets dominating. The main difference is that, within regions, the CpG triplets contribute complementary positive and negative peaks. In all three ancestral B allele scenarios, the class of site that should be most associated with introgressed fragments, A fixed in P2 (light blue trace), does not correlate with the overall signal and is small in magnitude, contributing only 8.5% of the total difference. Finally, the derived alleles (right hand panels) have lower overall counts and are dominated by complementary positive and negative values associated with sites where the A allele is fixed in either *P1* or *P2* (dark and light blue respectively). Here, none of the individual series correlate closely with the overall counts. Instead, in the numerically largest non-African – African comparison, the overall count shows two peaks, corresponding to triplets ACA, ACG and ACT, driven mainly by large differences at sites where B is fixed in P1 or segregating in *P1* and P2.

Although a focus on BABA – ABBA differences allows D to be broken down to expose the overall contribution of each class of site, it fails to capture information about the relative contribution per site. A small BABA – ABBA difference may actually reflect a large effect size if the number of sites in that class is relatively even smaller. Consequently, I replotted the data from the top left panel in Figure 1 for all classes of site (black line), replacing BABA – ABBA differences with D values (Figure 2). This plot contrasts with the equivalent profile in Figure 1 (black line), showing that the BABA – ABBA differences are little correlated with the number of sites. Interestingly, the XCG triplet, while still contributing high values, no longer stand out as unusual. Instead, the four triplets of the form XCC now appear unusal.

**Figure 2.**
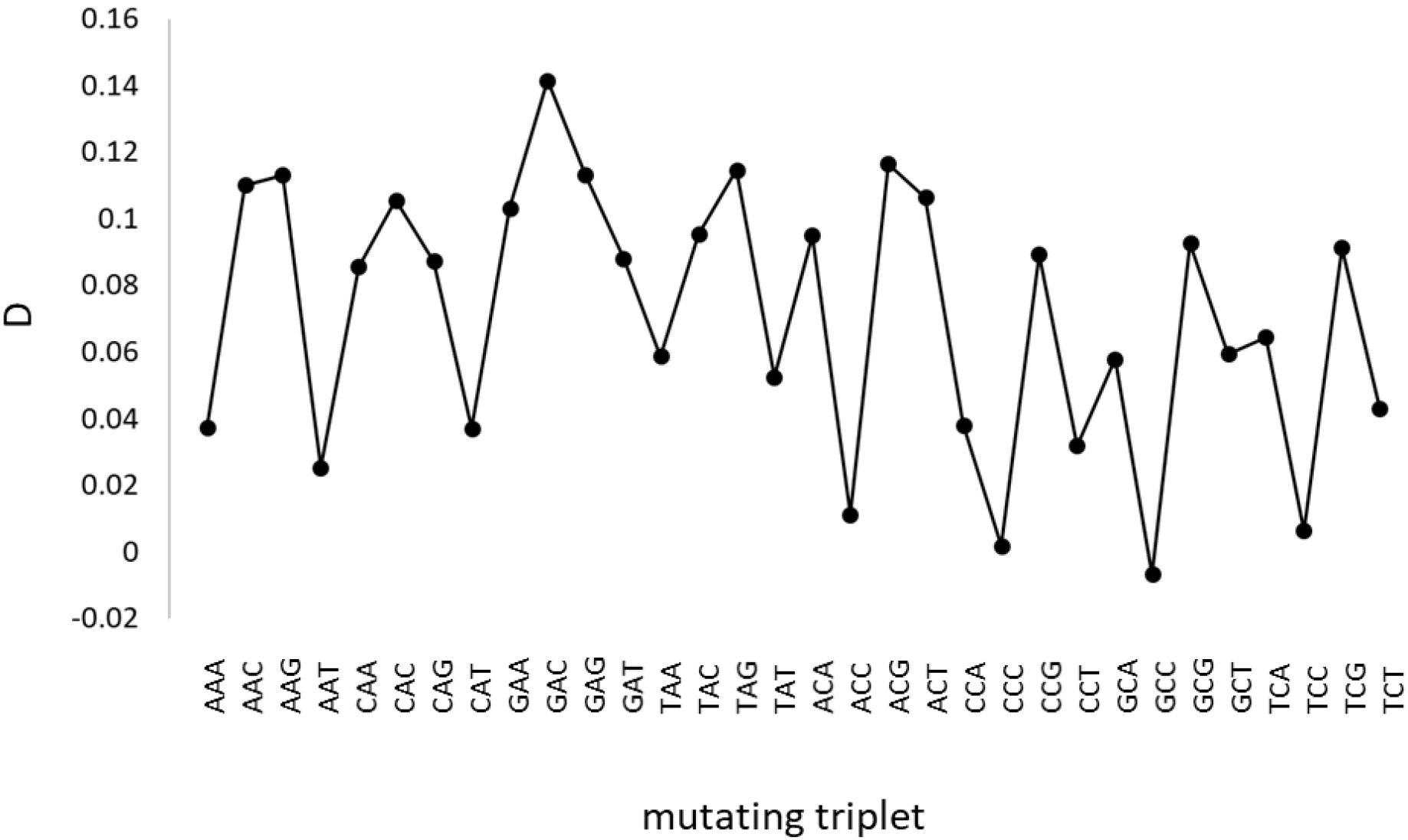
Variation in D with mutation type. Here, D was calculated separately for each of the 32 possible triplets for a standard non-African – African comparison, GBR-ESN. This plot is equivalent to combining the black line in the top two panels in Figure 1 and then dividing each value by its associated BABA + ABBA total count.

### b) Triplets, mutation rate and polymorphism

It is well known that CpG diads have very high mutation rates (25–27) and, as a result, are relatively rare across the genome. The over-representation of XCG triples in sites that contribute most to D therefore suggests that it is their mutation properties rather than introgression that drives the pattern. To understand this concept better, I counted triplet frequencies according to the status of the central base that mutated. To compare species, I counted sites where the base in one taxon (humans, Neanderthal, Denisovan or chimpanzee) differed from the other three. In each case, I counted the number of sites according to which of the 32 triplets formed the likely ancestral state, using only invariant sites in humans. I also explored the impact of polymorphism in humans and the possibility that parallel mutations on the chimpanzee lineage are likelier at sites with higher mutation rates. At polymorphic sites in humans I assumed that the ancestral state was indicated by the major allele, counted over the entire 1000 genomes data. Sites were sought where the ancestral human allele, Neanderthal and Denisovan bases are the same and then classified such sites according to: i) whether hominins and the chimpanzee differ; ii) whether humans are polymorphic or monomorphic. Summary triplet counts are presented in Figure 3, using the nomenclature [A/B]BBA to indicate sites where the human is polymorphic, Neanderthal = Denisovan = B and chimpanzee = A. As expected, the most striking feature involves XCG triplets, whose proportion varies 20-fold from 2.24 times the average frequency of all other triplets for [A/B]BBA to just 0.11 for AAAA (non-mutated sites). All other classes of site, including derived Neanderthal alleles (the Neanderthal base differs from both the chimpanzee and other hominins, black Xs) exhibit broadly similar profiles with a deficit of XCG triplets that is less extreme than for unmutated sites (AAAA, red).

**Figure 3.**
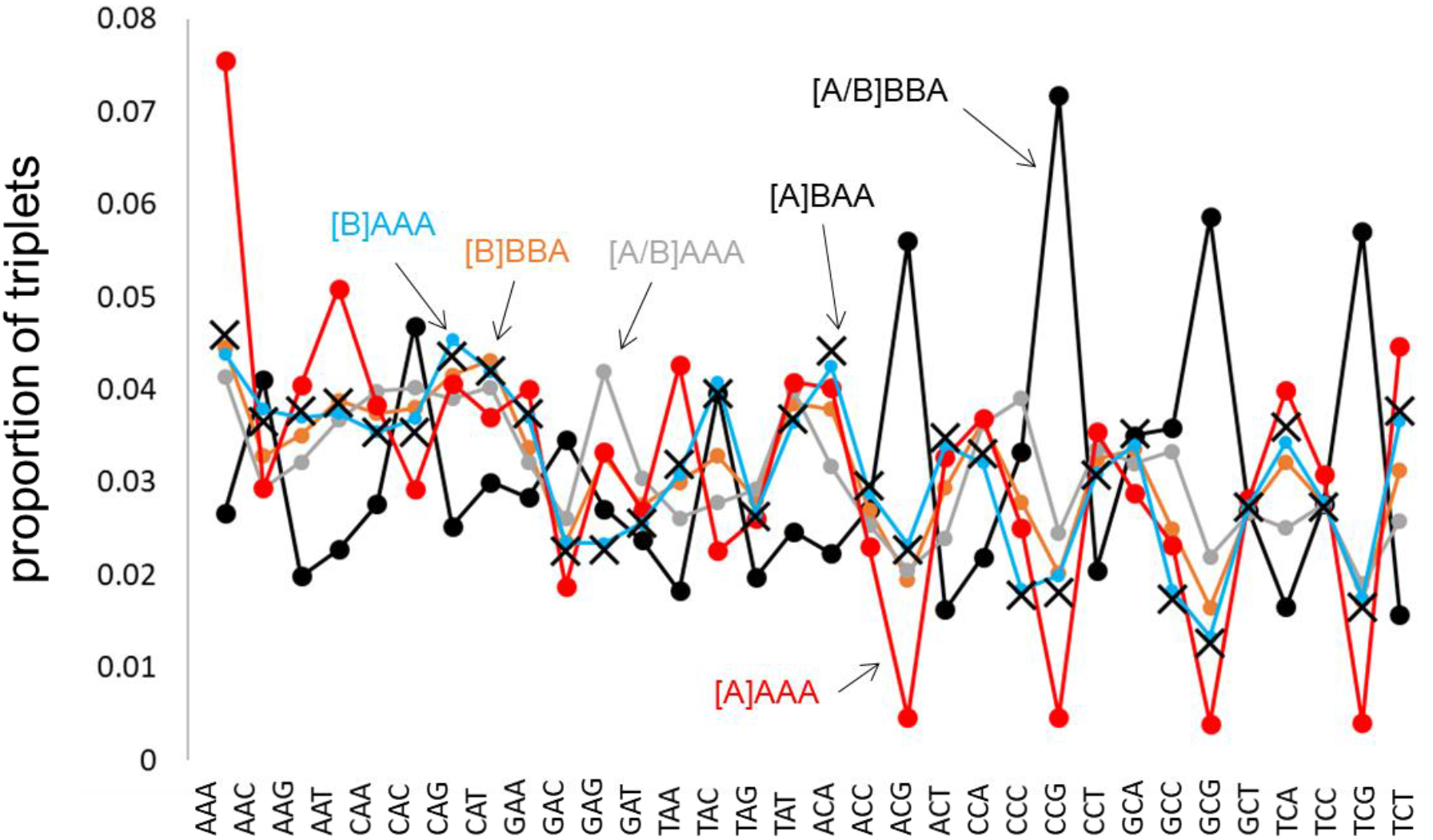
Frequencies of triplet base combinations in different classes of site. Autosomal genome alignments for humans, the Altai Neanderthal, the Denisovan and the chimpanzee where used to classify sites according to whether they are derived within a species, whether a substitution has occurred between hominins and the chimpanzee and whether the site is monomorphic or polymorphic in humans. Classifications are written [human]-Neanderthal-Denisovan-chimpanzee, with A/B indicating polymorphism. For each class, all qualifying triplets are counted and expressed as a proportion of all sites. Illustrated classes of site are: unmutated ([A]AAA, red); derived Neanderthal ([A]BAA, black crosses, noline); derived monomorphic bases in humans ([B]AAA, light blue); monomorphic in humans, derived in chimpanzees ([B]BBA, orange); polymorphic in humans ([A/B]AAA, grey); derived in chimpanzees, polymorphic in humans ([A/B]BBA, back). Connecting lines do not have a meaning and are added to aid visualisation only.

## Discussion

Here I uncover strong inter-relationships between D, a statistic used widely to quantify archaic introgression into humans, and characteristics of the substitution event that generated each informative polymorphism. Most strikingly, in the classic comparison between African and non-African humans, the signal captured by D is almost entirely attributable to sites where the putative introgressed allele is both ancestral to Neanderthals and Denisovans and is fixed outside Africa: removal of these sites reduces D to near-zero, 0.003. Moreover, the signal is dominated by sites where the mutating base is the highly mutable cytosine in CpG motifs.

Decomposing D reveals a striking pattern. The overall BABA – ABBA difference used to calculate ‘classical’ D (i.e. not decomposed) is 24,807 sites and is actually slightly lower than the difference contributed by sites where the Neanderthal B allele is fixed outside Africa and at high frequency inside Africa, 27,922. Restricting this class of site to those where the Neanderthal B allele is likely ancestral only reduces the difference to 23,731, still very close to the overall difference. Consequently, removal of these sites where B is fixed outside Africa reduces D to near- of just below zero, depending on whether the additional condition is applied. Unfortunately, in the classic model, where Neanderthals hybridised with humans somewhere around the Levant soon after the out of Africa event (28), most introgressed fragments will be absent from Africa and have low frequencies in non-Africans. This expectation is directly contradicted by the observed pattern. Moreover, the sites most likely to be generated by genuine introgression, those where the putative Neanderthal allele is both derived and restricted to non-Africans, contribute little to the overall signal and, perhaps tellingly, are essentially uncorrelated with the overall signal when the BABA – ABBA difference is decomposed according to which triplet mutated. Thus, in humans, Neanderthal introgression appears to contribute negligibly if at all to D as originally formulated.

A second major challenge for the introgression hypothesis is set by the dominant role played by sites containing highly mutable CpG triplets. Under a simple introgression model, D should only vary with triplet identity if the frequencies of different triplets differ appreciably between humans and Neanderthals. In reality, the frequencies are extremely similar. Thus, comparing triplet frequencies for derived Neanderthal alleles (i.e. [A][B][A][A]) with derived non-polymorphic human alleles, [B][A][A][A], the maximum difference in frequency seen between species at any one triplet is 10%, seen for CCG. Even allowing for the fact that a direct comparison is not possible due to lack of information about polymorphism in non-humans, 10% is clearly far too small to drive the ~400% excess signal seen at XCG triplets. Consequently, the XCG excess must be due largely to a process other than introgression.

D itself essentially captures information about the relative branch lengths of two human population / individuals to the test group, here the Neanderthal (1, 13). The branch to non-Africans is indisputably shorter and there appear only two ways this could be true, introgression into one population acting to mitigate divergence or a higher mutation rate in one lineage acting to increase divergence. Mutation rate variation is generally excluded through an explicit assumption (12, 13, 29), which means that any rejection of the null hypothesis must imply introgression. However, the current analysis shows that introgression contributes little if at all to the large positive values seen for D(*Europe, Africa, Neanderthal, chimpanzee*). By implication, the observed signal must be driven mainly by mutation rate variation. Here, the dominant role of XCG triples adds strong support. CpG is well known to be the most mutable diad (26), but the process is not reciprocal, in the sense that CG - > TG mutations occur at a far higher rate than TG - > CG. As a result, XCG triplets are generally rare compared with other motifs. Despite this, XCG triplets contribute almost 30% of the total signal captured by D, a dramatic excess that appears to reflect their greater mutability. Since most D-informative sites will be created by recurrent mutations that convert the common BBBA state (one substitution on the long branch separating hominins and chimpanzees), and the probability of two mutations at the same site scales with mutation rate squared, it is to be expected that the resulting signal is dominated by sites with unusually high mutation rates.

A model based on mutation rate variation is further supported by comparisons between populations from the same versus different major geographic regions. Within a major region (e.g. AFR-AFR or EUR-EAS), approximately symmetrical positive and negative BABA – ABBA difference peaks are seen for XCG sites where putative ancestral Neanderthal alleles are fixed in the first and second populations, respectively. This is consistent with the conditioning exposing equivalent B - > A mutations in both populations at similar rates. However, in the African – non-African comparison the negative peak, though present, is very much smaller than the positive peak. Such asymmetry would be the natural result of Africans having an appreciably higher mutation rate compared with non-Africans. It is this asymmetry that appears to be the primary driver of positive D values overall.

Finally, if D is driven mainly or even entirely by differences in mutation rate between populations, there remains the open question of why so many different studies report signals of introgression (3, 10, 30-32). Resolving this question should be the subject of future research. However, a key issue is that essentially all methods used to infer introgression are comparative, in the sense that they quantify how much more similar one human or one human population is to an archaic compared with a second. Asymmetry can arise either through introgressed fragments acting to increase similarity to the archaic or through a higher mutation rate acting to reduce similarity. The resulting patterns are not easy to distinguish. To do so requires careful attention to features like allele frequencies, which help inform whether it is the A or the B alleles driving any pattern. When this is done, the answer seems to become clearer, with most if not all of the signal being due to variable mutation rate.

In conclusion, partitioning D according to the characteristics of each site reveals a number of ways in which the data appear incompatible with D being driven by archaic legacies. At the same time, the data display several features that are both consistent with and indeed predicted by an alternative model in which variable mutation rates modulate the genetic distance between different human populations and related taxa such as Neanderthals. Surprisingly, the entire signal previously interpreted as being due to introgression appears instead to be driven by mutation rate differences. Such an apparently potent mechanism provides a simple general explanation for why the inference of introgression has become such a ubiquitous feature of hominin genetics.

## Methods

### Data

Data were downloaded from Phase 3 of the 1000 genomes project (33) as composite vcf files from (ftp://ftp.1000genomes.ebi.ac.uk/vol1/ftp/release/20130502/). These comprise low coverage genome sequences for 2504 individuals drawn from 26 modern human populations spread across five geographic regions: Europe (GBR, FIN, CEU, IBS, TSI); East Asia (CHB, CHS, CDX, KHV, JPT); Central Southern Asia (GIH, STU, ITU, PJL, BEB); Africa (LWK, ESN, MSL, GWD, YRI, ASW, ACB); and the Americas (MXL, CLM, PUR, PEL). Individual chromosome vcf files for the Altai Neanderthal genome were downloaded from http://cdna.eva.mpg.de/neandertal/altai/AltaiNeandertal/VCF/. Vcf files for the Denisovan genome were downloaded from http://cdna.eva.mpg.de/denisova/VCF/human/. I focused only on homozygote archaic bases, accepting only those with 10 or more reads, fewer than 250 reads and where >80% were of one particular base. This approach sacrifices modest numbers of (usually uninformative) heterozygous sites but benefits from avoiding ambiguities caused by coercing low counts into genotypes.

### Data Analysis

Analysis of the 1000g data and archaic genomes was conducted using custom scripts written in C++ (see Supplementary File 1 for the primary script). Other codes are available on request. Sites were only considered if a base was called in all taxa: humans, the Altai Neanderthal, Denisovan and chimpanzee. Since the 1000 genomes data include much imputation, individual genotypes are likely less reliable than population frequencies. Consequently, expected counts of ABBAs and BABAs were always calculated probabilistically based on population allele frequencies and assuming independent assortment. Mutating triplets were inferred from the human reference genome and converted to 32 possible type with either A or C as the central base, those with G or T being converted to their opposite strand equivalents. In all cases, the central base in the triplet that mutated was assumed to be the major allele in humans. This is an extremely close approximation because the overwhelming majority of variants are rare and are the product of relatively recent mutations.

## Supporting information

Example C++ code

ESM2 full output data

## Acknowledgements

I thank Simon Martin, Rob Foley and Marta Lahr for many useful discussions.

