## Supplementary material for "Decomposing the admixture statistic, D, suggests a negligible contribution due to archaic introgression into humans": Example C++ code

**C++ code used to calculate BABA – ABBA count differences in the 1000 genome Phase 3 data, partitioned according to the properties of the different site.**

This code should run if pasted into a compiler. I used Visual Studio 2019. Required input files will be uploaded to Dryad and comprise: ‘inpops.txt’, a file containing a list of 1000 genome samples, their population and region codes; and 22 individual chromosome files named ‘Master_CHR[#].txt’, each containing aligned bases for the Denisovan, Altai Neanderthal, chimpanzee, allele frequencies in each of the 26 human populations and flags describing the type of polymorphism.

```
// hangover headers, don't need all but was lazy and did not delete unneeded
#include <iostream>
#include <stdlib.h>
#include <fstream>
#include <string.h>
#include <stdio.h>
#include <math.h>
#include <time.h>

using namespace std;

void LoadPops();
void LoadReference(int chrom);
int triplet(long ln, char at);
void set_infile(int crm);
bool trans(char base1, char base2);
char infer(int b[4]);
int convert(char b);

char infile[200];
char line[1000000];
int REF[250000000]{};
int pops[2504][2]{};
int N_ind[26]{};
float ABBA[64][6][26][26][4]{};
long n = 0;
long long current = 0;


int main()
{
	// read population origin data for all 2504 individuals, stored as pops[ind][0 = pop, 1 = region]
	LoadPops();

   char temp[100000]; // recycled large character array

	// set and open output file
	char ofile[200] = "G:\\ABBATriplet.txt";
	ofstream out(ofile);
	if (out) cout << ofile << " is open\n";
	else cout << "output file NOT OPEN " << ofile, cin >> temp;

	// work through each chromosome in turn
	for (int chromosome = 1; chromosome < 23; chromosome++) {

		// load all bases for the reference genome for the current chromosome
		LoadReference(chromosome);

		// set and open the input file
		set_infile(chromosome);
		ifstream in(infile);
		if (in) cout << infile << " is open\n";
		else cout << infile << " is not open\n";

		// initialise a number of variables
		char cmp, nea, den;
		int SNP, MULT, IND, Nal;
		int prev = 0, lcn = 0, typ = 0;
		long locHUM = 0;

// work through the human data file, parsing the data line by line.  These are derivative
// master file containing aligned bases for Neanderthal, Denisovan and chimpanzee at each
// polymorphic human sites, human allele frequencies and flags for SNP / multiallelic / indel etc.
		while (in) {
			char alt[1000], ref[1000];
			in >> locHUM >> ref >> alt >> cmp >> nea >> den >> SNP >> MULT >> IND >> Nal;

			int freq[26][5]{};
			float fq[26][2]{};
			int NAL[3]{};
			for (int i = 0; i < 2; i++) {
				for (int pop = 0; pop < 26; pop++) {
					in >> freq[pop][i]; // allele counts
					fq[pop][i] = float(freq[pop][i]) / N_ind[pop]; // frequencies by population
					NAL[i] += freq[pop][i]; // overall allele counts
				}
			}
			in.getline(temp, 100000);

			lcn = locHUM / 1000000; // output progress in megabases
			if (lcn > prev) cout << prev << "\n";
			prev = lcn;

			// identify biallelic sites with all taxa scored and single base polymorpisms
			if (MULT==0 & Nal==2 & ref[1]=='\0' & alt[1]=='\0' & nea!='-' & cmp!='-' & den!='-' & nea!=cmp & SNP==1) {
				int trip = -1;
				int typ;

				// recover triplet value (0 - 31) for major human allele
				if (NAL[0] > NAL[1]) trip = triplet(locHUM, ref[0]);
				else trip = triplet(locHUM, alt[0]);

				// variable that reflects whether Neanderthal base is likely ancestral / derived
				if (nea == den) typ = 0;
				else typ = 2;

				// variable NE that for the Neanderthal based
				int NE = -1;
				if (nea == ref[0] & cmp == alt[0]) NE = 0;
				else if (nea == alt[0] & cmp == ref[0]) NE = 1;

				// if neand and chimp bases match ref and alt, calculate and count ABBAs and BABAs
				if (NE > -1) {
					for (int p1 = 0; p1 < 26; p1++) {
						for (int p2 = 0; p2 < 26; p2++) {
							// precalculate the probability of observing ABBA / BABA
							float ab = fq[p1][NE] * fq[p2][1 - NE];
							float ba = fq[p1][1 - NE] * fq[p2][NE];

	// store ABBAs and BABAs according to criteria: 0 = all sites, 1 = sites where Neanderthal all
   // is fixed in population 1, 2 = non-Nea fixed in pop 1, 3 = Nea fixed in pop 2, 4 = non-Nea
   // fixed in pop 2 and 5 = polymorphic in both populaitons (else).

ABBA[trip][0][p1][p2][0 + typ] += ab, ABBA[trip][0][p1][p2][1 + typ] += ba;
if (fq[p1][NE] == 1) ABBA[trip][1][p1][p2][0 + typ] += ab, ABBA[trip][1][p1][p2][1 + typ] += ba;
else if (fq[p1][1-NE]==1) ABBA[trip][2][p1][p2][0 + typ] += ab, ABBA[trip][2][p1][p2][1 + typ] += ba;else if (fq[p2][NE]==1) ABBA[trip][3][p1][p2][0 + typ] += ab, ABBA[trip][3][p1][p2][1 + typ] += ba;
else if (fq[p2][1- NE]=1) ABBA[trip][4][p1][p2][0 + typ] += ab, ABBA[trip][4][p1][p2][1 + typ] += ba;
else ABBA[trip][5][p1][p2][0 + typ] += ab, ABBA[trip][5][p1][p2][1 + typ] += ba;
						}
					}
				}
			}
		}
		in.close();
		in.clear();
	}

	// output the ABBA and BABA counts for each class
	for (int block = 0; block < 6; block++) {
		for (int p1 = 0; p1 < 26; p1++) {
			for (int p2 = 0; p2 < 26; p2++) {
				out << block << "\t" << p1 << "\t" << p2;
				for (int trip = 0; trip < 32; trip++) {
					out << "\t" << ABBA[trip][block][p1][p2][0] << "\t" << ABBA[trip][block][p1][p2][1];
					out << "\t" << ABBA[trip][block][p1][p2][2] << "\t" << ABBA[trip][block][p1][p2][3];
				}
				out << "\n";
			}
		}
		out << "\n";
	}

	out.close();
}

// convert location and major allele into triplet number, 0 – 31, 32 = no triplet returned
int triplet(long ln, char at)
{
	int mid = -1;
	if (at == 'A') mid = 1;
	else if (at == 'C') mid = 2;
	else if (at == 'G') mid = 3;
	else if (at == 'T') mid = 4;
	if (REF[ln - 1] * mid * REF[ln + 1] > 0) {
		if (REF[ln] > 2) {
			return (5 - mid - 1) * 16 + (5 - REF[ln + 1] - 1) * 4 + 5 - REF[ln - 1] - 1;
		}
		else {
			return (mid - 1) * 16 + (REF[ln - 1] - 1) * 4 + REF[ln + 1] - 1;
		}
	}
	return 32;
}

void set_infile(int crm)
{
	char tmp[5];
	strcpy_s(infile, "G:\\Master_CHR");
	if (crm < 10) tmp[0] = crm + 48, tmp[1] = '\0';
	else tmp[0] = (crm / 10) + 48, tmp[1] = crm - (crm / 10) * 10 + 48, tmp[2] = '\0';
	strcat_s(infile, tmp);
	strcat_s(infile, ".txt");
}

// transition / transversion test.  Can be used to restrict classes further
bool trans(char base1, char base2)
{
	if ((base1 == 'A' & base2 == 'G') | (base2 == 'A' & base1 == 'G')) return false;
	if ((base1 == 'C' & base2 == 'T') | (base2 == 'C' & base1 == 'T')) return false;
	return true;
}

void LoadPops() // read in population and region codes for each of 2504 individuals
{
	char infile1[200] = "B:\\inpops.txt";
	ifstream in1(infile1);

	in1.getline(line, 1000);
	for (int i = 0; i < 2504; i++) {
		in1 >> line;
		in1 >> pops[i][0];
		in1 >> pops[i][1];
		pops[i][0]--; // population codes start at 1 so decrement for use as array indices
		pops[i][1]--;
		N_ind[pops[i][0]] += 2; // counts alleles scored in each population
		in1.getline(line, 200);
	}
	in1.close();
}

void LoadReference(int chrom)
{
	char infile2[200] = "B:\\hs37d5.fa";
	ifstream in2(infile2);
	if (!in2) cout << "not open " << infile2 << "\n";
	// in2.seekg(current);

	int cr = 0;
	while (!in2.eof() && cr < chrom) {
		in2.getline(line, 1000);
		if (line[0] != 'A' && line[0] != 'C' && line[0] != 'G' && line[0] != 'N' && line[0] != 'T') {
			cout << line << "\n";
			cr++;
		}
	}

	long cnt = 0;
	long c = 0;
	char base = 'A';
	while (!in2.eof() && cnt < 249999999 & base != '>') {
		base = in2.get();

		// store upper case bases A, C, G, T and N as numbers
		if (int(base) > 64 && int(base) < 87) {
			cnt++;
			c++;
			if (c > 1000000) {
				cout << cnt << "\n";
				c = 0;
			}
			if (base == 'A') REF[cnt] = 1;
			else if (base == 'C') REF[cnt] = 2;
			else if (base == 'G') REF[cnt] = 3;
			else if (base == 'T') REF[cnt] = 4;
			else REF[cnt] = 0;
		}

		// handle rare lower case bases
		else if (int(base) > 97 & int(base) < 118) {
			cout << ".";
			cnt++;
			c++;
			if (c > 1000000) {
				cout << cnt << "\n";
				c = 0;
			}
			if (base == 'a') REF[cnt] = 1;
			else if (base == 'c') REF[cnt] = 2;
			else if (base == 'g') REF[cnt] = 3;
			else if (base == 't') REF[cnt] = 4;
			else REF[cnt] = 0;

		}
	}
	in2.close();
	in2.clear();
}
```
